# SpotLink enables sensitive and precise identification of site non-specific cross-links at the proteome scale

**DOI:** 10.1101/2021.12.31.474672

**Authors:** Weijie Zhang, Pengyun Gong, Yichu Shan, Lili Zhao, Hongke Hu, Qiushi Wei, Zhen Liang, Chao Liu, Lihua Zhang, Yukui Zhang

## Abstract

We developed SpotLink software for identifying site non-specific cross-links at the proteome scale. Contributed by the dual pointer dynamic pruning (DPDP) algorithm and the quality control of cross-linking sites, SpotLink identified more than 3000 cross-links from human proteome database with rich site information in a few days. We demonstrated that SpotLink outperformed other approaches in terms of sensitivity and precision on a simulated dataset and a protein complexes dataset with known structures. Additionally, we discovered some valuable protein-protein interaction (PPI) information contained in the protein complexes dataset and HeLa dataset, indicating the unique identification advantages of site non-specific cross-linking. The excellent performance of SpotLink will increase the usage of site non-specific cross-linking in the near future. SpotLink is publicly available on GitHub [https://github.com/DICP1810/SpotLink].

## Introduction

Chemical cross-linking combined with mass spectrometry (CXMS) has emerged as a critical method for protein structure construction and protein-protein interaction (PPI) analysis^1-5^. To date, a variety of cross-linkers have been developed for different applications. Among them, the most widely used cross-linkers are those with amino-reactive groups (lysine or N-terminal of peptides), such as DSS, BS3, and DSSO due to their high reactivity^6^. However, these cross-linkers may not be suitable for some circumstances. Firstly, biological systems such as membrane proteins or proteins associated with the nervous system may be deficient in lysine^7^. Nearly 46% of human proteins in the UniProt database have lysine ratios that are less than the average ratio **(Supplementary Fig. 1)**. Secondly, lysine-restricted cross-linking limits the coverage of distance information, which is necessary for modeling proteins^5, 8^. To overcome these issues, cross-linkers with high amino acid reactivity, such as succinimidyl 4,4’-azipentanoate (SDA) and formaldehyde, have been developed. Furthermore, these site non-specific cross-linkers have been successfully used to analyze some biosamples and obtain general interaction information regarding target proteins^9-11^.

However, few attempts have been made to investigate protein structures and interactions at the proteome-wide level by utilizing site non-specific cross-linkers due to the complexity of the data analysis process^12^.

The first challenge encountered when interpreting site non-specific CXMS data is the expansion of the computational search space. In site-specific cross-linking, the possible peptide pairs of one spectrum among N peptides is *O (N*^*2*^)^13^. However, as for site non-specific cross-linking, if one of the reactive groups of the cross-linker is site non-specific, the number of combination candidates expands from *O (N*^*2*^) to *O (MN*^*2*^), where M denotes the average number of reactive peptide residues. If the cross-linkers reactive groups are all site non-specific, the site combinations of two peptides will be quadratic and reach *O (M*^*2*^*N*^*2*^). Currently available software tools, such as xi^14^, pLink2^15^, Kojak^16^, and MeroX^17, 18^, are built for site-specific cross-linking identification and are inadequate for identifying cross-links in such a vast space. The existing tools attempt to support this process with two strategies. MeroX tries to limit the search space to one or two cross-linking sites, which causes the loss of many potential site combinations, resulting in decreased software sensitivity. Xi tries to make all amino acids available for searching, which exacerbate performance issues while identifying proteome-wide site non-specific cross-links. Thus, a search space with a complexity of *O (MN*^*2*^) or *O (M*^*2*^*N*^*2*^) becomes a new performance bottleneck for site non-specific CXMS data interpretation.

The second challenge is the quality control difficulty induced by the different cross-linking sites combinations obtained from a large space. For tools designed for site-specific cross-link identification, it is assumed that the correct residue pairs are usually limited to a few amino acids, such as lysine-lysine, and these residue pairs share the same confidence level^19^. However, this assumption does not hold for site non-specific cross-linking because of the existence of sites competition, which causes site specific software to report some site combinations that are unlikely to occur in biological samples when identifying site non-specific cross-links, increasing the risk of reporting false-positive results. Although the current software designs multilevel quality control workflows to guarantee the validity of reports, including those at the spectrum level, peptide level, and protein level, it still cannot guarantee the precision of cross-linking site location under the competition of different site combinations^20-22^. Hence, software that comprehensively considers cross-linking sites scoring is required, for site non-specific cross-link identification.

In this work, we built SpotLink software to identify site non-specific cross-linking peptides at a proteome scale with identification efficiency and cross-linking sites localization effectiveness **(Supplementary Fig. 2**). Firstly, an enumeration-based search strategy was induced to recall potential peptide pairs on each spectrum. SpotLink, unlike fragment index-based software, attempts to cover the entire peptide area and site combination space with *O (MN*^*2*^) or *O (M*^*2*^*N*^*2*^). The dual pointer dynamic pruning (DPDP) algorithm was designed to report the top-20 coarse-scoring candidates with a balance between speed and sensitivity. With the benefit of the DPDP algorithm and efficient memory operations, SpotLink performs as swiftly as index-based strategy software such as pLink2 and xi, hence enabling the identification of site non-specific CXMS data at the proteome scale.

Secondly, to ensure quality control regarding different site combinations for site non-specific cross-linking, SpotLink was designed with an algorithm to calculate the site cross-linking probability within each cross-link for scoring and a ranking algorithm to estimate the site false discovery rate (sFDR) from a global view. By splitting the probability distributions of positive and negative site combination samples, SpotLink could exclude random site combinations with the probability score predicted by the sFDR. The sFDR provided a comprehensive evaluation of various site combinations and create a statistic model for the quality control of site non-specific cross-linking residue pair; this approach is simply adaptable to any site non-specific cross-linker.

Finally, to demonstrate the performance and robustness of SpotLink, we analyzed four site non-specific cross-linking datasets, including simulated data, the amyloid beta (Aβ) aggregate data, the condensin complex data, and human cell data. On the simulated dataset, SpotLink achieved up to a 10% improvement in cross-linking site identification precision over existing software. On the protein and protein complex dataset, SpotLink identified 30% more cross-linking sites with high confidence, thus achieving the highest cross-linking coverage for identifying site non-specific CXMS data. On the largest human cell dataset, SpotLink identified 3494 cross-links from the human proteome in ∼10 days with strong evidence. These results show that SpotLink is capable of identifying site non-specific data ranging from simple protein assembles to whole proteome proteins.

## Results

As a general overview, we first introduced the algorithmic advances of SpotLink in identifying site non-specific cross-links. Then, we created a simulated SDA cross-linking dataset to demonstrate the necessity of developing site non-specific cross-link identification software. We benchmarked SpotLink with two popular software programs, pLink2 and xi. As a result, it was determined that site-specific tools are insufficient for identifying site non-specific cross-links. Furthermore, by analyzing the Aβ dataset and the previously published condensin complex dataset, we proved that site non-specific cross-linking coupled with SpotLink provided more structural information about proteins and protein complexes. In addition, the Aβ dataset represents an example of studying low lysine proteins with site non-specific CXMS. Finally, SpotLink was able to identify site non-specific cross-links at the human proteome scale in days with high confidence. We performed a comprehensive analysis on the HeLa membrane dataset.

### Workflow of SpotLink

SpotLink consists of four parts in its backend workflow: the preprocessing step, the searching step, the scoring step, and the quality control step. As a brief introduction, SpotLink takes protein sequence files and mgf files as inputs (**Fig. 1a**). The protein sequence files are prepared into a peptide list, and the spectra from mgf files are preprocessed; this step is followed by recalling the candidate peptide pairs with the DPDP algorithm. After the recalling step, the top 20 candidate peptides for each spectrum are kept for the quality control step. In the quality control step, SpotLink conducts reranking at the peptide level and calculates the false discovery rate (FDR) as all software does. In addition, SpotLink uniquely scores and calculates the sFDR scores. The results obtained at the peptide level and the site level are given separately.

**Fig. 1.**
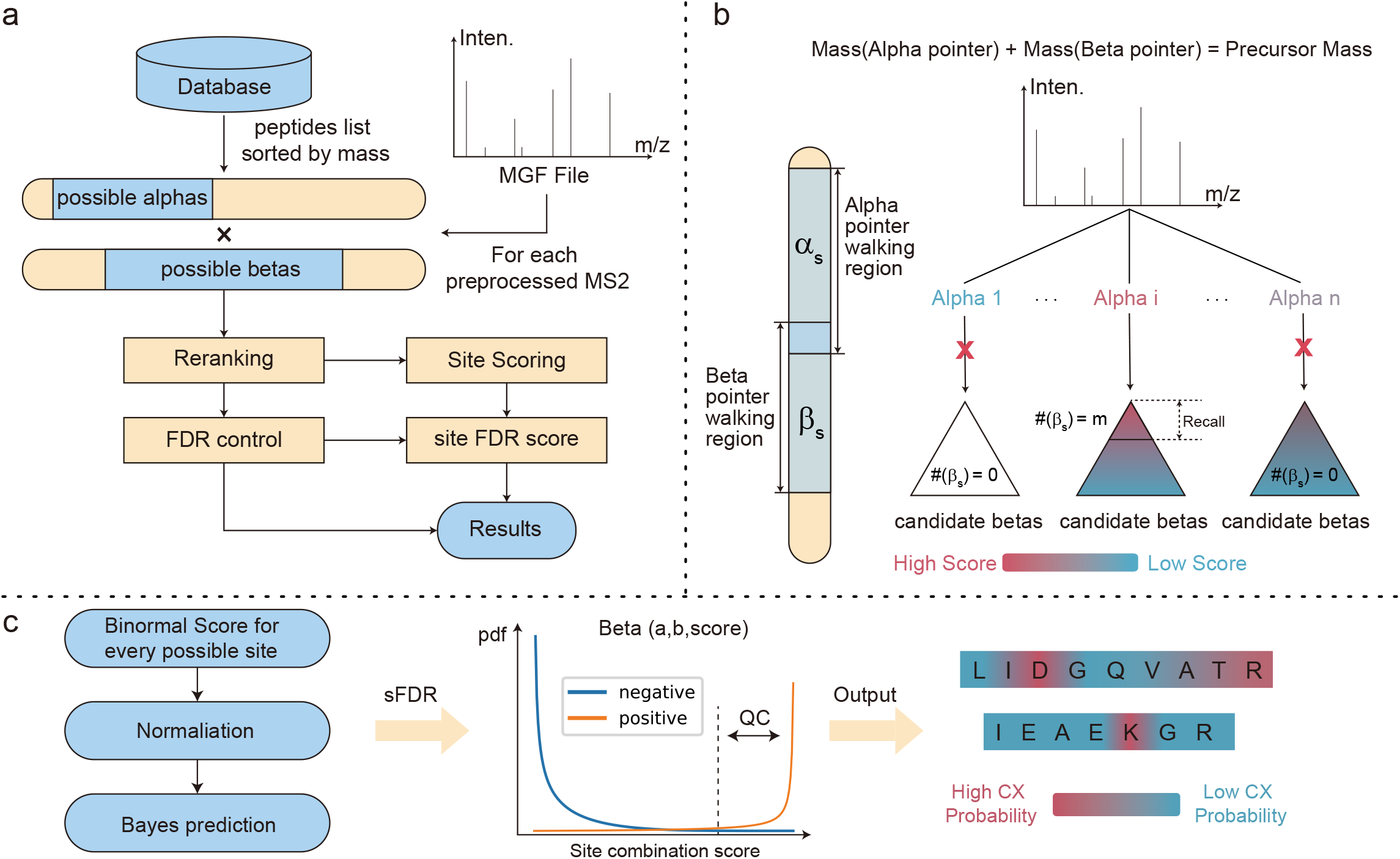
Algorithm workflow of SpotLink. (**a**) Overview. Step 1, protein sequences and MGF files are preprocessed for subsequent use. Step 2, for each spectrum, enumerated searching is performed to recall the top 20 cross-linking peptide pairs. Step 3, quality control on peptide level, including FDR calculation for each peptide. Step 4, quality control on cross-linking site level, including site scoring and site FDR (sFDR) score calculation. (**b**) The sub-workflow of enumerated searching. To reduce redundant matches, we designed the Dual Pointers Dynamic Pruning (DPDP) algorithm. Dual pointers can reduce redundant pairwise matching and scoring, besides dynamic pruning can determine whether to perform in-depth for the β peptides given a α peptide. n represents the number of total α peptides. m represents the number of recall β peptides. (**c**) The sub-workflow of site quality control. This step contains multiple levels of scoring. For each spectrum: (1) Perform binomial distribution scoring on all possible cross-linking sites. (2) Calculate the normalized probability from the site score. (3) Predict the reliability based on the Bayesian model. For all results: (4) Sort and report high confidence cross-linking sites based on the Beta distribution of positive and negative samples.

The DPDP algorithm (**Fig. 1b**) consists of two walking pointers and a scissor for tree data structures. The two pointers include an α-peptide pointer with a greater mass and a β-peptide pointer with a lower mass. For each spectrum, the two pointers walk along with the peptide list with a precursor mass restriction. All combinations form a candidate peptide pair tree. The tree pruning behavior of the algorithm triggered under two conditions. Firstly, if the α-peptide score falls below the threshold, the β peptides that match the α peptide are pruned. Secondly, the combination of pointers is pruned if the combined α-peptide and β-peptide score is less than the threshold or if the two scores cannot form a suitable cross-linking site pair. By employing this algorithm, it is possible to ensure that the program recalls the proper candidate sequence within a reasonable amount of time. The quality control module for cross-linking sites contains multiple steps (**Fig. 1c)**. Firstly, the binormal scores of different site combinations are utilized to evaluate the reliability of each peptide spectrum match (PSM). The score values provide confidence levels for each cross-linking site within each PSM. Secondly, the sFDR of each cross-linking pair is calculated based on the expectation–maximization (EM) algorithm with the Beta distribution. The sFDR score indicates the degree of mismatching between each cross-linking pair in a global context, and a lower score is better. SpotLink can minimize the probability of reporting low-confidence cross-linking site combinations by implementing these tactics.

### Evaluation of SpotLink using the simulated dataset

To test the compatibility of existing software on identifying site non-specific cross-linking, the simulated SDA cross-linking dataset was created. We performed ten experiments. Each contained 2,000 spectra generated from 300 randomly selected *E*.*coli* proteins **(Methods)**. All spectra were complete containing only the full sets of b and y ions generated by the two cross-linking peptides, and they did not contain any extraneous noise or interfering peaks **(Supplementary Fig. 3**). The software was intended to report correct results for all spectra. We performed searches on the limited 300 proteins used to create the spectra as well as all *E*.*coli* proteins. In pLink2 and xi, the cross-linking sites were set to all amino acids. We compared the sensitivity, and precision values at the peptide level and cross-linking site level, as well as the time consumptions of these tools **(Fig. 2**).

**Fig. 2.**
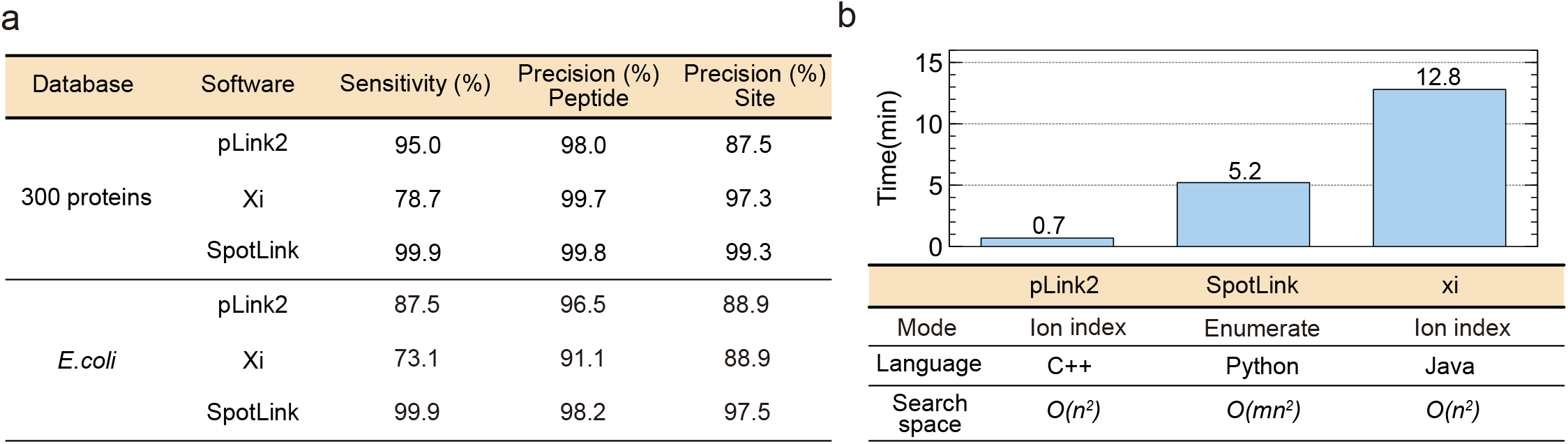
Performance on the simulated SDA cross-linking dataset. (**a**) Ten repetitions were carried out to obtain the average value of sentitivity, precision on peptide, and precision on cross-linking site. In each repetiton, the simulated-SDA dataset was searched among pLink2, xi, SpotLink using two databases, one consists of 300 proteins, and the other contains all proteins from *E*.*coli*. FDR was set at 5%. (**b**) The average time-consuming of three software searching on the *E*.*coli* proteome. And the differences among search strategy, programming language, search space were shown.

Although pLink and xi had great precision at the peptide level, their precision in terms of cross-linking site location was indeed underestimated under the same peptide FDR of 5%. As a result, they were unreliable in identifying site non-specific cross-linking sites. Under the same FDR (5%) conditions, SpotLink obtained the highest sensitivity, peptide precision, and cross-linking site precision **(Fig. 2**). The analysis showed that SpotLink achieved impressive performance both on limited proteins and the proteome, which was attributed to the DPDP algorithm and the site level quality control mechanism. Additionally, we compared the search times of the three software on the *E*.*coli* proteome. SpotLink, which was coded in Python as enumeration-based software with a large searching space, had the same time overhead order of magnitude as the index-based software.

Although the simulated data were created under idealized conditions and were simpler than real-world data, only SpotLink could achieve accuracies greater than 97% in terms of both aspects. Its outstanding performance provided a solid foundation for the subsequent analysis of more samples.

### Validation and new insight into Aβ aggregates and the condensin complex

Site non-specific cross-linking provides more site information regarding the structures and interactions of proteins and protein complexes. We analyzed two datasets, the Aβ protein aggregate dataset and the previously published condensin complex dataset, to demonstrate this fact.

Aβ, a small protein containing only two lysine residues in its sequence, is aggregated in different forms, and is a major neuropathological hallmark of Alzheimer’s disease (AD), which is not suitable to be researched by lysine-lysine cross-linking^23, 24^. Aβ monomers can form higher-order assemblies ranging from low molecular weight to high molecular weight oligomers, protofibril, fibrils, and senile plaques. We gathered these aggregates and cross-linked them by SDA to create the Aβ dataset.

SpotLink reported 32 unique cross-link sites pair from the Aβ dataset at a 5% peptide FDR **(Fig. 3a)**. All of these results had sFDR score less than 1e-5. Among them, a number of 14 interactions contacted with the Lys (16), 16 interactions contacted with the Lys (28), and one interaction contacted with the N-terminal of Aβ. We mapped our cross-linking data onto the published electron microscopy (EM) structure of the dimer aggregates (PDB entry 6SHS) and the solid-state NMR structure of the trimer aggregates (PDB entry 2M4J) **(Fig. 3b)**. We set the maximum Cα-Cα distance between the residues that SDA could cross-link to 30 Å^10^. For the dimeric aggregate, 30 (out of 32) Cα-Cα distances were less than 30 Å, and the maximum mapping Cα-Cα distance is 35.3 Å. For the trimeric aggregates, 31 (out of 32) Cα-Cα distances were less than 30 Å, and the maximum mapping Cα-Cα distance is 40.9 Å **(Supplementary Fig. 4a, b**).

**Fig. 3.**
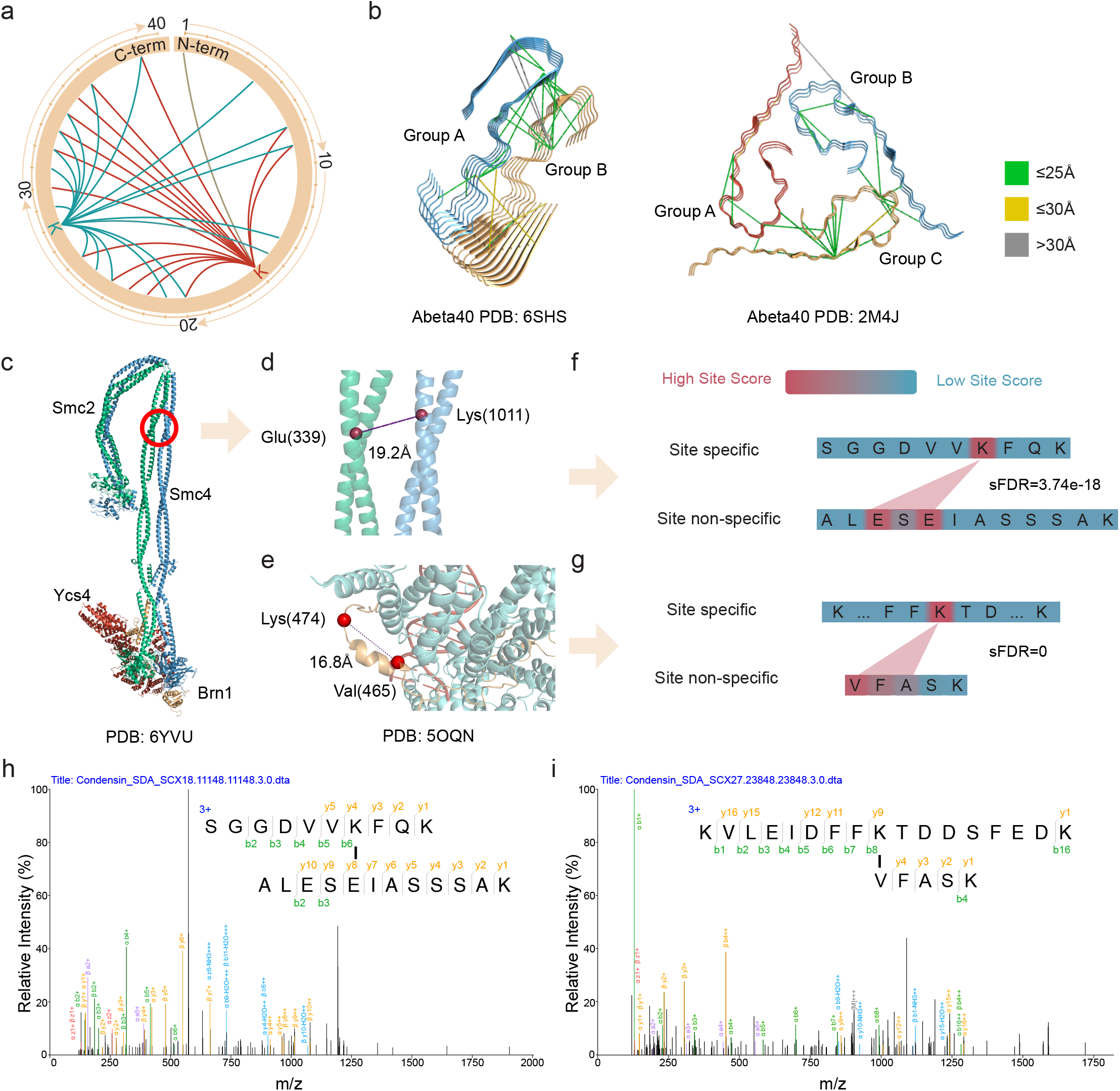
SpotLink result for the Aβ dataset and the condensin dataset produced by site non-specific cross-linking. (**a**) The SpotLink identification results from Abeta dataset under 5% PSM FDR. In the PPI network, each line in the circle represents a potential cross-linking site pair. (**b**) We then matched the cross-linking information on two PDB structures 6SHS and 2M4J, which represent the dimeric and the trimeric of Abeta aggregates respectively. Most of the cross-linked pairs match the structure under 30 Å. (**c**) The structure of the condensin complex. Two cross-linking site pairs identified by SpotLink uniquely were shown here. (**d**) Glu (Smc2, 339) – Lys (Smc4, 1011) was mapped to structure (PDB ID: 6YVU) with 19.2 Å. (**e**) Lys (Brn1, 474) – Val (Brn1, 465) was mapped to structured (PDB ID: 5OQN). (**f**) The sFDR score from SpotLink of these two cross-linking pairs. (**g**) Tandem mass spectrum of Glu (Smc2, 339)-Lys (Smc4, 1011) interprotein cross-linking. (**h**) Tandem mass spectrum of Lys (Brn1, 474) – Val (Brn1, 465) intraprotein cross-linking.

As one of the highlights, different Aβ aggregates could have distinct structural characteristics, and site non-specific cross-linking combined with SpotLink could exactly report these cross-linking sites as expected. Three cross-linking site pairs were reasonable only on one formation of Aβ aggregate. **(Supplementary Fig. 4c)**. Residues 8 and, 9-28 (Gly, Ser-Lys) were well matched with the trimeric aggregated Aβ, and residues 1-16 (Asp, Lys) were well matched on the dimeric aggregated Aβ. These differences could not be obtained via site-specific cross-linking. By analyzing this dataset with SpotLink, we confirmed the outstanding ability of site non-specific cross-linking and gained some fresh insight into the aggregation of Aβ.

To illustrate the benefits of site non-specific cross-linking in the study of protein complexes, we chose the condensin dataset, which has become the newest milestone in the application of site non-specific cross-linking in biological systems. The condensin complex consists of five proteins: Smc2, Smc4, Brn1, Ycs4, and Ycg1. The recently published EM structure (PDB entry 6YVU) clearly analyzed its structure **(Fig. 3c)**. We downloaded this dataset and performed an analysis with SpotLink.

By reanalyzing the data, SpotLink reported a ∼30% increase in the number of cross-linking sites over that in the original report at 1% FDR **(Supplementary Fig. 5a, b)**. We visualized the interactions between these five subunits that were reported with at least two supporting PSMs **(Supplementary Fig. 5c)**. Clearly, SpotLink achieved excellent cross-linking coverage on this dataset. By mapping these results to existing and AlphaFold predicted structures^25^, we evaluated the cross-linking sites pair reported by SpotLink **(Supplementary Fig. 6a, b)**. More than 80% of the cross-links could be mapped to the corresponding structures within 30 Å. Interactions that could not be verified by their structures could be supported by other evidence including PSMs.

To further demonstrate the accuracy of SpotLink identification results, we chose two cross-links reported only by SpotLink and demonstrated their reliability from three aspects: spectrum matching, cross-linking site scoring, and structural matching. The interprotein cross-linking between Smc2(Glu, 339) and Smc4(Lys, 1011) could be identified by peptides SGGVVKFQK and ALESEIASSSAK **(Fig. 3h)**. The sites score heatmap shows that site non-specific cross-linking happened among E(338)S(339)E(340) from Smc2 subunit; And SpotLink reported the cross-links with an sFDR score 3.74e-18, indicating that the randomness of cross-linking between these residues was low **(Fig. 3f)**. Besides, this cross-linking pair could be mapped on the 6YVU structure with 19.2 Å **(Fig. 3d)**. The intraprotein cross-linking between Brn1(Lys, 474) and Brn1(Val, 465) could be identified by the peptides KVLEIDFFKTDDSFEDK and VFASK **(Fig. 3i)**. The sites score heatmap shows that site non-specific cross-linking happened among V(465)F(466)A(467); And SpotLink reported the cross-links with an sFDR score 0, which implies a high level of trust **(Fig. 3g)**. Besides, this cross-linking pair could be mapped on the 5OQN structure with 16.8 Å **(Fig. 3e)**. These findings demonstrate effectiveness and robustness of SpotLink while exploring site non-specific cross-linked protein complexes.

### Proteome-wide profiling of in-vivo cross-linking from HeLa samples

The algorithmic improvement of the SpotLink algorithm allowed it to perform site non-specific cross-links identification against a complete human proteome database. We prepared in-vivo cross-linked Hela cells by SDA and isolated membrane-related cell components for the following mass spectrometry analysis.

SpotLink reported 3647 cross-linked PSMs and 3494 cross-linking sites at 1% FDR **(Fig. 4a, Supplementary Data)**. To test the validity of our data, we verified the results of SpotLink report in terms of both the matching structure and the protein interaction network. Firstly, we mapped our cross-linking sites to 399 high-resolution structures yielding distance information for 473 intraprotein interactions **(Supplementary Data)**. By measuring the probability distribution of these distance information, it was clear that the majority of cross-links were within 30 Å and were significantly distributed at approximately 10 Å **(Fig. 4b)**.

**Fig. 4.**
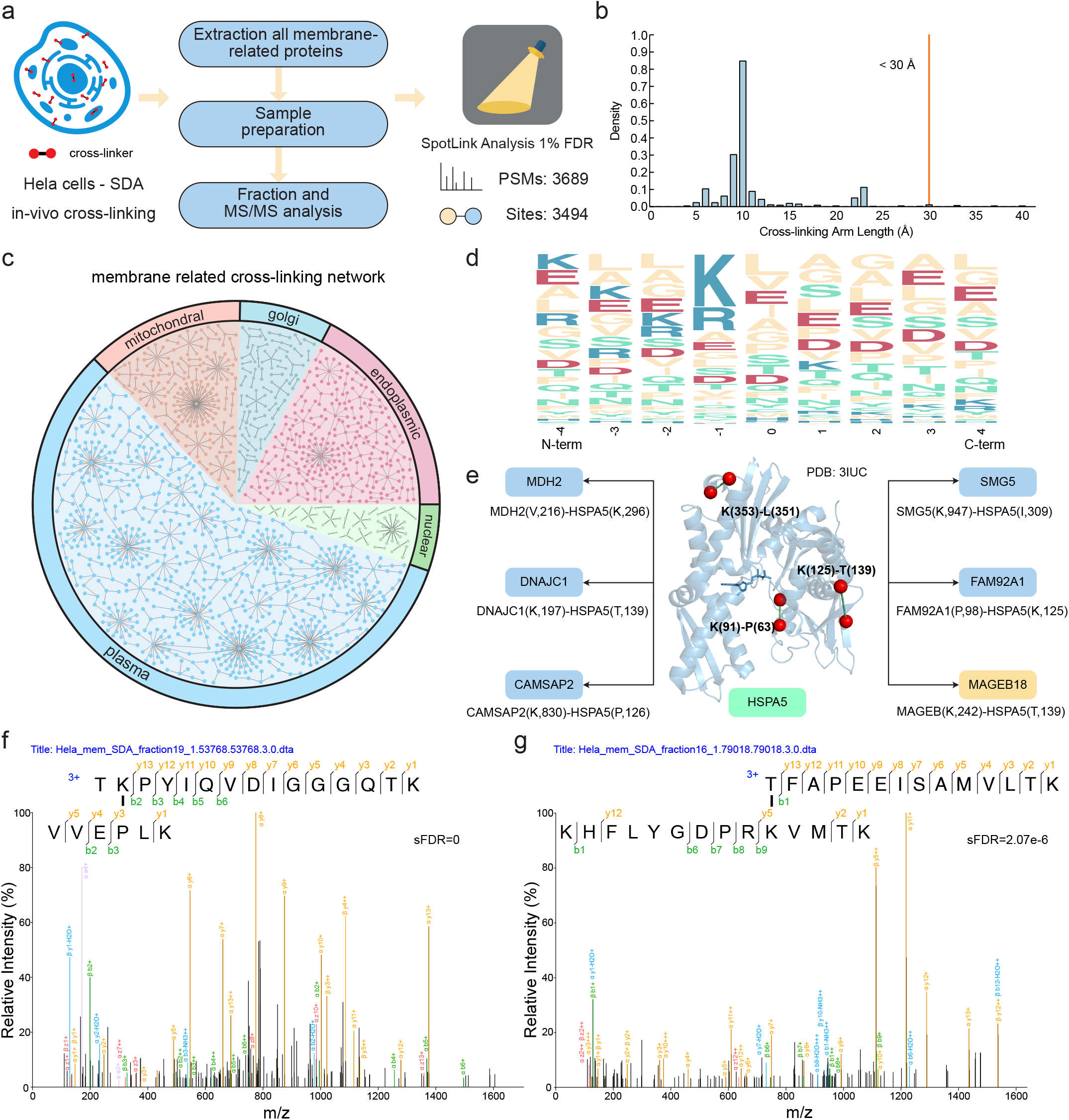
SpotLink results of the Hela dataset from the human proteome scale. (**a**) The general description of the Hela dataset and Spotlink analysis results at 1% FDR. (**b**) The cross-links arm length distribution. The result is generated by mapping 399 PDB structures with SpoLink results. Most of the arm length located within 30 Å. (**c**) The PPIs networks of each membrane-related cell component clustered by cellular location. The spots represent identified gene. Each line between the spots represents an identified cross-link. And the area of each subcellular membrane component sector represents the number of PPIs. Please refer to supporting for more details. (**d**) The amino acids frequency analysis around the cross-linking site from the non-specific reaction terminal of the cross-linker SDA at the proteome scale. The size of the letters represents the frequency. Anionic amino acids are marked in red, and cationic amino acids are marked in blue. For other amino acids, polar amino acids are marked in green, and non-polar amino acids are marked in yellow. **e**) Examples of PPIs of HSPA5 identified by SpotLink. The structure of HSPA5 encoded protein with PDB structure 3IUC mapped the K(353) - L(351) with 6.6 Å, the K(91) -P(63) with 6.8 Å, the K(125)-T(139) with 11.6 Å. The PPIs in the blue chunk were confirmed by the STRING database, and the PPI in the yellow chunk was newly identified by SpotLink. (**f**) Tandem mass spectrum of Lys (HSPA5,125) - Pro (FAM92A1, 98) interprotein cross-linking. (**g**) Tandem mass spectrum of Lys (HSPA5, 242) - Thr (MAGEB18, 139) interprotein cross-linking.

Secondly, to illustrate how the proteins connected and to understand the content of the membrane related PPIs, we generated a PPI network and categorized these PPIs into five main compartmental networks (i.e., the plasma membrane, endoplasmic reticulum membrane, mitochondrial inner membrane, Golgi membrane, and nuclear membrane) **(Fig. 4c, Supplementary Fig.7)**. The numbers of interprotein interactions contained in these interaction networks were 764, 196, 168, 75, and 63, respectively **(Supplementary Data)**. In these networks, many structurally well-studied proteins with suitable distance mappings, PPIs with database support, and interactions with strong PSM support could be observed **(Supplementary Fig. 8, 10)**. One example was the HSPA5 encoded protein P11021, which is an endoplasmic reticulum chaperone that plays a key role in protein folding and quality control in the endoplasmic reticulum lumen. Our data provided evidence regarding direct intraprotein and interprotein interactions in the cellular context around P11021 **(Fig. 4e)**. All the intra cross-links, K(353)-L(351), K(125)-T(139), and K(91)-P(63), from P11021, could be well matched to the protein structure 3IUC (solved by X-ray). And several inter cross-links could be confirmed by spectra matches and STRING (search tool for the retrieval of interacting genes/proteins) supports, such as the PPIs between HSPA5 with MDH2, DNAJC1, CAMSAP2, SMG5, and FAM92A1. Besides, SpotLink provided new information about previously unknown contacts of HSPA5. For example, the cross-linking between HSPA5 with MAGEB18 had solid correspondence with a high PSM score, suggesting a novel protein interaction **(Fig. 4g)**.

Furthermore, to study the precision values and tendencies of the SpotLink reported cross-linking sites at the proteome scale, we performed a statistical analysis of the cross-linking sites on this scale. By calculating the frequencies of the sFDR scores for all possible cross-linking sites, it was clear that the sFDR scores of most dependable cross-linking sites were approximately zero, while a large number of random cross-linking sites had sFDR scores that were located near 1 **(Supplementary Fig. 9a)**. In other words, the lower the sFDR score given by SpotLink was, the more reliable the cross-linking site was at a proteome-wide scale. For instance, the cross-linking between HSPA5(Lys, 125) and FAM92A1(Pro, 98) was identified by peptides TKPYIQVDIGGGQTK and VVEPLK with an sFDR score 0 **(Fig. 4f and, Supplementary Fig. 9b)**, and the cross-linking between HSPA5(Thr, 139) and MAGEB18(Lys,242) could be identified by peptides TFAPEEISAMVLTK and KHFLYGDPRKVMTK with an sFDR score 2.07e-6 **(Fig. 4g and, Supplementary Fig. 9b)**. These results were credible from cross-linking site scale. Finally, we analyzed the frequencies of amino acids from the site non-specific cross-linking peptides that had sFDR scores less than 0.05 as reported by SpotLink **(Fig. 4d)**. According to the results of the motif analysis, few amino acids, including D, E, K, R, L, A, G, and S, expressed higher affinity to SDA in terms of site non-specific cross-linking at the human proteome-scale; these results were similar to prior findings obtained at the protein scale ^26^.

## Discussion

To tackle the barrier facing site non-specific cross-linking identifying, we developed SpotLink, which enabled multi-sites, high-coverage, and high-accuracy interaction investigation at the proteome scale. Thus, the benefits of site non-specific cross-linking for exploring a wider range of protein systems with a greater theoretical coverage could be fully utilized. We primarily made two algorithmic advances. Firstly, we developed the DPDP algorithm to recall cross-links with sensitivity and balanced time-consuming. The algorithm enabled SpotLink to report cross-links with high sensitivity. Secondly, we designed the algorithm for quality control of cross-linking sites, which provided confidence scores to each site pair locally (site binormal score) and globally (sFDR) and enabled SpotLink to achieve high cross-linking site precision. Furthermore, a large number of software engineering optimizations make SpotLink a user-friendly and robust research platform.

By testing SpotLink with four different levels of datasets: the simulated dataset, Aβ dataset, condensin complex dataset, and HeLa dataset, we demonstrated a breakthrough in identifying site non-specific CXMS data. SpotLink exhibited excellent sensitivity, precision, time-consuming, and cross-linking coverage at different scales and provided new insights into protein interactions and structures with high confidence by considering different cross-linking site combinations.

Site non-specific cross-linking combined SpotLink has been successfully used to identify different samples, but there is still the potential to improve and achieve higher cross-linking coverage. The future compatibility of SpotLink for identifying cleavable cross-links will significantly benefit the study of site non-specific cleavable cross-links, especially formaldehyde cross-links, which are links with high reactivity and high permeability into cells and tissues. In addition, site-independent cross-linking based on the data-independent acquisition (DIA) method and the corresponding identification algorithm in SpotLink is expected to improve the identification of cross-linking sites. More importantly, site non-specific cross-linkers with enrichment groups are expected to be designed. Increased peptide abundance, as a result of enrichment strategies, will enables site non-specific cross-linking to yield higher analysis depth when combined with SpotLink.

